# Synergistic effects of hypergravity and immune challenge on the zebrafish larval transcriptome

**DOI:** 10.64898/2026.09.30.755701

**Authors:** Maria Mitsopoulou, Eloi Casals, Anna Esteve-Codina, Laia Ribas

## Abstract

The transcriptomic interaction between altered gravity and innate immunity remains largely unexplored. In this study, we established a novel dual-stress model combining 3*g* hypergravity with lipopolysaccharide (LPS) immune challenge in 5 days post-fertilization (dpf) zebrafish (*Danio rerio*) larvae. RNA sequencing revealed that exposure to hypergravity alone induced no differentially expressed genes (DEGs), indicating a remarkable transcriptional resilience. In contrast, the addition of an immune challenge under hypergravity conditions triggered a substantial transcriptomic response involving 2,706 DEGs. Functional enrichment analyses identified significant alterations in pathways related to immune function, development, and circadian regulation. Targeted qPCR analyses further validated the expression patterns of selected genes. These findings suggest that while zebrafish larvae can effectively tolerate hypergravity alone, the combination of gravitational and immune stressors elicits profound molecular responses. This study provides new insights into the interplay between altered gravity and innate immunity using an animal model, contributing to future strategies aimed at safeguarding astronaut health and supporting space-based aquaculture systems.

## Introduction

Nowadays, humanity has entered the new era of space exploration. As a result, it is essential to understand the biological response to altered gravitational fields to ensure the safety of future long-term space missions. Gravity is a fundamental and constant physical force that has shaped the evolution of all living organisms on Earth ^1^. Although spaceflight primarily exposes organisms to microgravity, hypergravity conditions provide valuable ground-based experimental models. Hypergravity is defined as the conditions in which gravity exceeds Earth’s levels (1*g*) ^2^ and helps to decipher how gravitational changes influence physiological and molecular mechanisms ^3,4^. Research into hypergravity conditions typically involves centrifugation devices with sufficient radius to generate inertial forces within the biological samples^5^. In general, changes in gravitational load influence a broad spectrum of biological processes, including bone development, cardiovascular function, and immune responses^4^. In fact, hypergravity has been documented to modulate immune activity across diverse taxa, ranging from insects ^6^ and amphibians ^7^ to mammals ^5^ and teleost fish ^8^. In addition, living systems possess specialized sensory mechanisms to perceive and respond to gravity. In vertebrates, the vestibular system is the principal organ for maintaining equilibrium and coordinating movement. In fish, exposure to altered gravity disrupts vestibular function, leading to orientation challenges and physiological stress^9^.

Animal models are useful to comprehend the impact of different factors of long-term space flights on living organisms and can be helpful forecasting negative effects of space flights on humans ^10^. Zebrafish (*Danio rerio*) is one of the most important world-wide vertebrate model organisms in developmental biology, genetics, neurophysiology, biomedicine and for aquaculture research ^11–13^. It has a short life cycle, rapid development, and high genomic similarity with humans. It is easy to maintain, it has well-characterized physiology and its embryos are transparent. Fish are considered ideal models for gravitational studies because their aquatic environment provides a condition of near-neutral buoyancy during development. Among fish species, zebrafish have become an essential vertebrate model for gravity-related research due to their experimental advantages and biological relevance ^11,14^. However, previous studies exposing zebrafish embryos to hypergravity reported significant physiological and morphological impacts, such as body curvature, tail and body deformation ^15^. Behavioral studies indicated that hypergravity induces erratic swimming behavior, jerky movements, and a shift towards a vertical ascendant position, due to environmental stress ^15^. In addition, hypergravity affects otolith —utilized for orientation— early development in zebrafish leading to overgrowth as well as it significantly affected the musculoskeletal system^9,16^. Moreover, exposure between 3 and 5 dpf altered the mechanical material properties of jaw cartilage and the composition of the extracellular matrix (ECM). Similarly, 3*g* exposure from 5 to 9 dpf led to a significantly larger head skeleton and increased bone formation in specific cranial bones^17^.

Despite these insights, there is limited knowledge on molecular and physiological interactions during space flights ^18^. Evidence from both human astronauts and animal models consistently demonstrates that exposure to the space environment induces significant alterations in immune function ^19–21^. In humans, investigations on astronauts that returned from either short or long-term International Space Station (ISS) missions revealed a significant dysregulation of immunological function, including a 20–85% suppression of natural killer cell cytotoxicity and a decrease in T-lymphocyte proliferative activity ^21^. The space environment therefore caused immunodeficiency, which was associated with a higher clinical risk of infections and autoimmune disorder ^22^.

While the effects of spaceflight on immune function have been documented in humans, considerably less is known about the molecular mechanisms underlying these responses in aquatic vertebrates. Studies performed in fish models, both under real and simulated spaceflight conditions on Earth, revealed important immune-related molecular responses. ISS studies on fish model organisms such as Japanese medaka (*Oryzias latipes*) identified the intestine as a highly space-responsive tissue by transcriptomic analyses showing the upregulation of over 2,500 genes. These changes included the modulation of immune-related pathways across multiple tissues, suggesting immune vulnerability and stress-related responses under spaceflight conditions ^20^. Moreover, under simulated spaceflight conditions, zebrafish embryos exposed to continuous gamma radiation exhibited significant transcriptomic shifts in key developmental and immune-related pathways ^23^. Overall, the use of these aquatic models provides essential insights into the immune and physiological adaptations required to ensure the safety of both biological support systems and human crews during long-duration space missions ^24^.

Although previous studies have demonstrated that space-associated stressors can induce significant immune-related transcriptional changes in fish models, how gravitational stress influences immune responses at the transcriptomic level remains poorly understood. To address this gap, the present study expands previous hypergravity research by introducing an immune stimulant, lipopolysaccharide (LPS). LPS, a major component of the outer membrane of Gram-negative bacteria, triggers a strong innate immune response without inducing systemic pathology ^25^. Given the high degree of conservation of immune regulatory pathways between zebrafish and mammals, LPS has become a widely used model for investigating acute inflammatory responses in fish ^26^.

The primary aim of this study was to investigate, for the first time, the synergistic effects of hypergravity and immunological challenge on zebrafish from a transcriptomic perspective. To achieve this, the research was structured around three core objectives. First, we seek to determine the optimal immune challenge dose eliciting a measurable response without compromising viability. Second, using RNA-sequencing strategy, we aimed to identify differentially expressed genes (DEGs) which serve as transcriptomic biomarkers for hypergravity-induced stress and immune activation. Finally, a subset of immune-related genes identified by RNA-seq was selected for expression validation, thereby confirming the robustness of the transcriptomic findings.

## Materials and Methods

### Zebrafish husbandry and experimental groups

AB strain adult zebrafish were maintained at the Experimental Aquarium Zone (ZAE) at the Institute of Marine Sciences (ICM-CSIC, Barcelona, Spain). The animals were housed in 9 L tanks into a recirculating system (Aquaneering, San Diego, CA). Environmental conditions within the chamber were strictly controlled, including a 12-hour light/dark photo period, an air temperature of 26 ± 1°C, and a humidity of 60 ± 3%. Water quality was managed through a 3,000 L/h pump and a UV sterilization system to ensure the elimination of bacterial pathogens. Physicochemical parameters were recorded daily to ensure stability: water temperature was kept at 28 ± 0.2°C, pH at 7.2 ± 0.5, conductivity between 750–900 μS, and dissolved oxygen at 6.5–7.0 mg/L ^12,13^. Additionally, commercial kits were used for the weekly monitoring of ammonia, nitrate, sulfate, and sulfite levels. The diet of adult fish was consisted of twice-daily feedings including both dried food and live *Artemia nauplii* (AF48, INVE Aquaculture, Dendermonde, Belgium).

Single zebrafish pairs were breed overnight inside the breeding tanks. Upon removal of the divider the following morning, fertilized eggs were collected, counted and maintained in Embryo Medium solution (EM, pH = 7.2; NaCl 0.8 g/L, KCl 0.04 g/L, NaH_2_PO_4_ 0.0036 g/L, H_2_KO_4_P 0.006 g/L, CaCl_2_ 0.144 g/L, MgSO_4_ 0.12 g/L, NaHCO_3_ 0.35 g/L, and 20 μl/L Trypan blue as antifungal). Fertility was verified based on standard reference values for this species and post-hatch survival based on the Fish Sexual Development Test guidelines (OECD, 2011).

### Generation of hypergravity conditions

Hypergravity conditions were generated based on an existing design ^15^ using a custom-made centrifuge constructed from a standard laboratory rotary mixer (model ANR100DE, OVAN laboratory equipment) capable of reaching 100 rpm. This design features two perpendicular arms (R = 25 cm) terminating in opposing gondolas designed to support experimental 96-well plates, covered with an adhesive film. To ensure a stable environment for the larvae, the centrifuge was enclosed in a methacrylate chamber equipped with a heating system managed via ProClima v8.1.6.1 software (Schneider Electric 2020). The device was operated at its maximum velocity of 100 rpm (10.47 rad/s), producing a centripetal acceleration (a_c_) of 27.4 m/s^2^. The a_c_ added to the existing Earth acceleration (*g*) of 9.81 m/s^2^ resulted to a total acceleration (a_T_) of 29.1 m/s^2^, which is equivalent to 2.96 g (~3*g*).

### LPS dose-response for immune stimulation

To establish a robust model for immunological challenges under gravitational conditions, dose-response survival assays were conducted. Fertilized eggs were transferred individually into 96-well plates containing 250 µL of Embryo Medium. A total of 200 embryos from three biological pairs were used. The assay was divided into five LPS concentrations, allocating 40 embryos per dose, which were distributed equally across four experimental conditions (n=10 per group). The groups were defined as: (1) Control (CT): samples maintained under standard Earth gravity (1*g*), (2) Hypergravity (HG): samples subjected to 3*g* hypergravity, (3) LPS: immune stimulated samples with no gravitational changes (1*g*), and (4) LPSHG: samples stimulated with LPS and subjected to 3*g* hypergravity.

To determine the optimal, non-lethal dose for inducing an immune response while maintaining the survival levels for long-term observation, the following concentrations were used: 1 μg/ml, 10 μg/ml, 15 μg/ml, 20 μg/ml, 40 μg/ml. Within each concentration, only the LPS and LPSHG groups were exposed to the designated LPS concentration. The CT and HG groups were maintained in standard EM. Survival rates were monitored daily from 0 to 5 dpf and compared across all four groups. The hypergravity device was stopped for less than five minutes each day for these observations.

### RNA extraction and RNA sequencing

Following five days of experimental exposure, zebrafish larvae were euthanized via thermal shock (cooling) for five minutes and following preserved at –80°C for subsequent analysis. Total RNA was extracted using the Quick-RNA Microprep Kit by ZymoResearch based on a standardized protocol. The procedure began with sample homogenization, performed using 300 ml of lysis buffer. Following extraction, the quality and quantity of RNA were assessed using the NanoDrop Spectrophotometer which measures concentration (ng/ml) and chemical purity via 260/280 and 260/230 absorbance rations and the TapeStation which provides the RNA Integrity Number (RIN). Only samples demonstrating high integrity (RIN > 8.0) were selected for RNA sequencing. The total RNA was sent to the National Center for Genomic Analysis (CNAG, Barcelona, Spain) for high-throughput short-read RNA sequencing which was executed on a total of four representative samples for each group, except for the LPSHG group in which one sample failed the library preparation. RNA samples libraries were prepared with the KAPA mRNA HyperPrep Kit and sequenced on an Illumina NovaSeq 6000 (2×151 bp).

### Bioinformatic analysis

RNA-seq reads were mapped against the *Danio rerio* genome (GRCz11) using STAR 2.7.8a ^28^ with ENCODE parameters. Transcript quantification was performed with RSEM 1.3.0 ^29^ using the Ensembl 110 (GRCz11.110) annotation. Genes with at least 1 count-per-million reads (cpm) in at least 4 samples were retained. Counts were normalized using the trimmed mean of M-values (TMM) method and transformed into log2-counts per million (logCPM). Differential gene expression analysis was performed using the limma R package v3.54.2 ^30^. The voom function ^31^ was used to estimate the mean-variance relationship and compute observation-level weights. These voom-transformed counts were used to fit linear models. Functional enrichment analysis was conducted using g:Profiler via the gprofiler2 R package v0.1.8 ^32^. Multidimensional scaling plots were generated with the limma function plotMDS using the top 500 most variable genes. Gene set enrichment analysis was performed using FGSEA v1.24.0 ^33^ with Reactome pathways for *Danio rerio*, using a preranked list based on the moderated *t*-statistic from Limma. Raw sequencing data was submitted to the National Center for Biotechnology Information (NCBI) with the GEO accession number: GSE334374. (To review GEO accession GSE334374: Go to https://www.ncbi.nlm.nih.gov/geo/query/acc.cgi?acc=GSE334374. Enter token uhcrmswubfejdah into the box).

### Functional analysis

Initially, DEGs were evaluated across all possible pairs between experimental groups to assess the independent impacts of hypergravity and immunostimulation. To interpret the broader biological significance of these transcriptomic shifts, a Gene Ontology (GO) enrichment analysis was conducted.

For the LPSHG vs CT comparison, targeted filtering was applied to the DEGs using an absolute log2-Fold Change (|Log_2FC| > 2) and an adjusted p-value < 0.03. From this dataset, the DEGs were ranked by absolute log2FC in descending order and any uncharacterized genes were excluded. Then, candidates that overlapped with the top 10 enriched GO categories were identified. From these overlapping genes, individual functions were verified using ZFIN, Ensembl (GRCz12tu), and relevant literature, ultimately selecting the top 10 genes related to immune system functions and the top 10 genes related to developmental processes.

Conversely, for the LPSHG vs LPS comparison, we focused on the top three enriched categories and selected the overlapping DEGs without further filtering. Subsequently, to isolate the core genetic drivers of the combined hypergravity and immune response, the overlap of these functionally curated DEGs between the two most biologically relevant comparisons (LPSHG vs LPS and LPSHG vs CT) was evaluated using a Venn Diagram ^34^.

### Validation through quantitative PCR

To validate the RNA-seq findings, the dataset of genes was filtered (adj. p-value < 0.05 and Absolute log2FC > 1.5) to ensure robust validation targets. Following these criteria, 11 genes from the two comparisons were isolated for further experimental validation.

Primers were designed based on the coding DNA sequence (CDS) of the selected zebrafish genes using the NCBI database (**Table 1**). The optimal primers were restricted to a product size of 70-120 bp, a melting temperature (T_m_) of 61.5 ± 1,5 °C, and GC content >50%. The qPCR validation was performed to ensure primer stability and efficiency. Regarding the efficiency standards, primers were required to meet an R^2^ > 0.985, a slope between −3.1 and −3.6 and an efficiency between 90 and 110%.

**Table 1.**
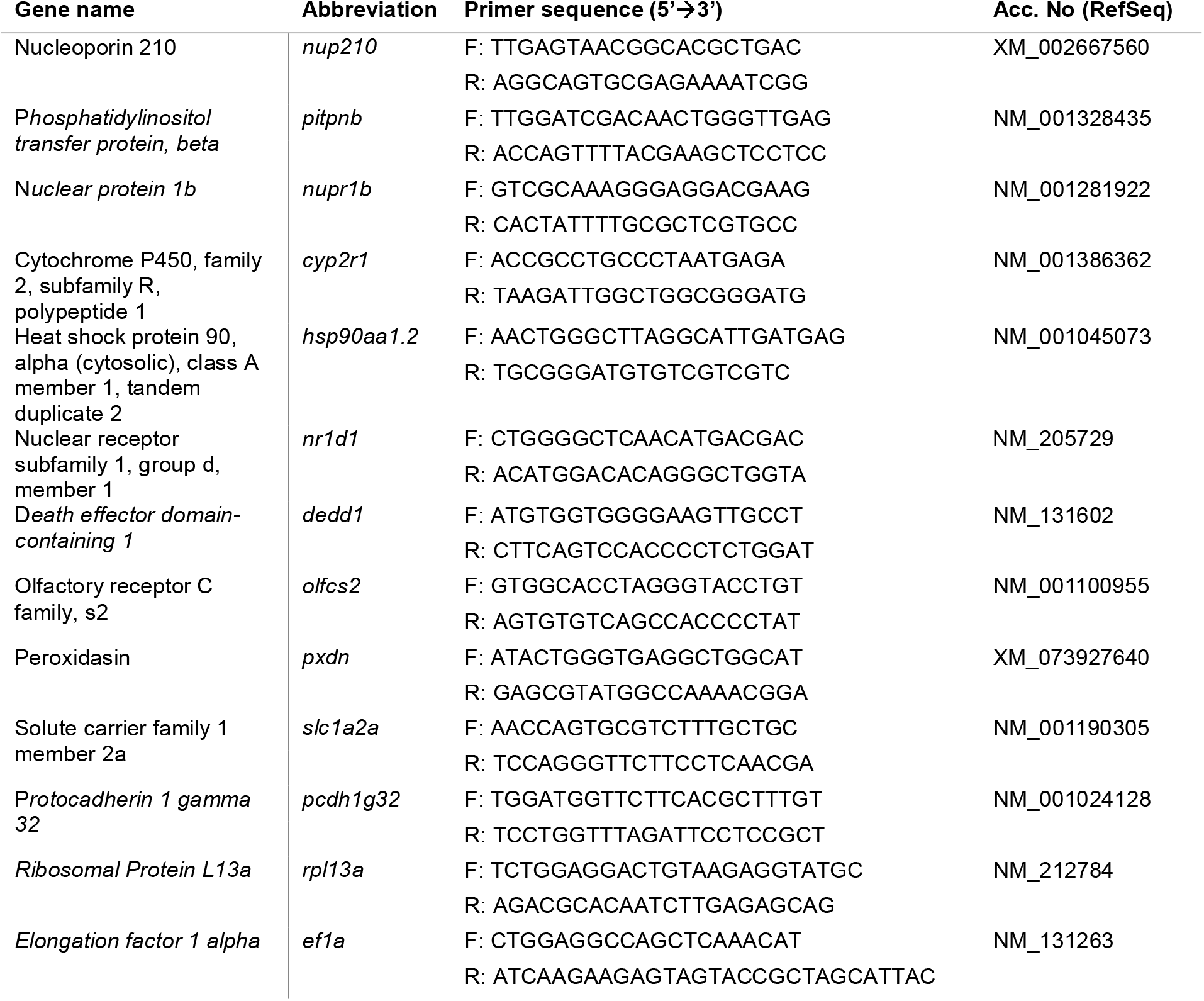
Gene specific primers used for RNA-seq validation by quantitative PCR.

Total RNA of each sample was reverse-transcribed into cDNA using the NZY First-Strand cDNA synthesis Kit (Nzytech) according to the manufacturer’s instructions. Real-time amplification was performed in a 384-well format using SYBR Green. Each 10μL reaction volume consisted of 1μL of sample cDNA, 5mL SYBR Green, 3μL Nuclease Free H_2_0 and 0.5μL of each Forward and Reverse primer. Data was normalized using housekeeping genes *rpl13a* and *ef1a*. Finally, log2FoldChange data from qPCR were compared against the RNA-seq log2FoldChange data to verify consistent biological trends.

### Ethics statement

All zebrafish procedures were performed in accordance with the guidelines approved by the Bioethical Committee of the ICM. ICM fish facilities are officially validated for animal experimentation by the Ministry of Agriculture and Fisheries (REGA no. ES080190036532).

## Results

### Determination of optimal LPS dosage for immunological challenge in zebrafish larvae

The survival assays revealed a clear dose-dependent relationship between LPS concentration and gravitational force (**Figure 1**). At lower concentrations (1 μg/ml to 10 μg/ml), survival remained above 70% throughout the 5-day observation period. However, a significant shift towards mortality was observed at the higher concentrations of LPS. The 15 and 20 μg/ml doses showed a moderate reduction in viability, with survival dropping to 70% by 5 dpf for the LPSHG group. The highest concentration (40 μg/ml) was highly toxic for the LPS and LPSHG groups, in which mortality rates exceeded 80% by 2 dpf. This concentration was considered unsuitable.

**Figure 1.**
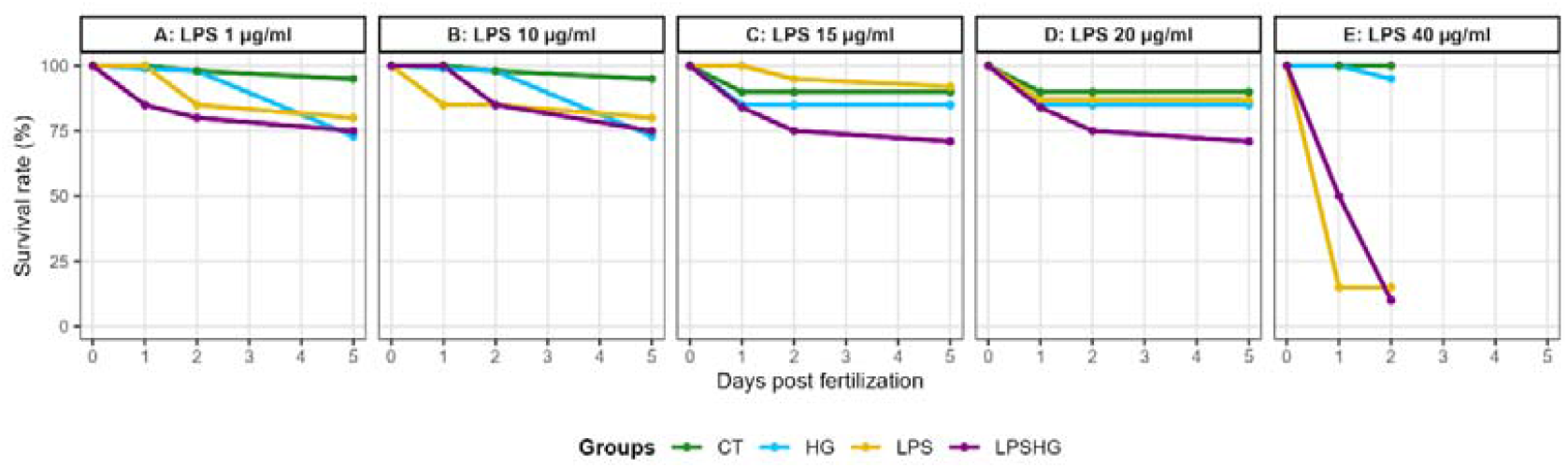
Survival rates of zebrafish larvae following exposure to varying concentrations of lipopolysaccharide (LPS). Zebrafish embryos were exposed to LPS concentrations of 1, 10, 15, 20, and 40 μg/ml at 0 days post-fertilization (dpf). Survival was monitored daily over a 5-day period to determine the optimal non-lethal dose for immunological challenge. Graphs A–E represent dose-response curves compared across the four different experimental groups: CT, HG, LPS and LPSHG. A clear dose-dependent decrease in survival is observed, with 40 μg/ml being lethal by 2 dpf for the LPS and LPSHG groups, while doses between 1 and 10 μg/ml maintained high survival rates suitable for long-term observation.

Across all the tested doses, the CT and HG groups maintained high survival rates (often >90%), confirming that any mortality observed during these experiments was due to the treatments. Based on these data, concentrations between 1 to 10 μg/ml were identified as optimal as they provided a sufficient challenge to the immune system while maintaining a robust survival of larvae for long-term observation.

### Transcriptomic response to dual stressors

To evaluate the molecular impact of hypergravity and immune stimulation, a comparative transcriptomic analysis was performed across all experimental groups. For the LPS and LPSHG groups, samples with a 1 μg/ml LPS concentration were used in the transcriptomic analysis as this represented the lowest dose capable of inducing detectable immune alterations. The distribution of DEGs, classified as upregulated and downregulated, is summarized in **Table 2**.

**Table 2.**
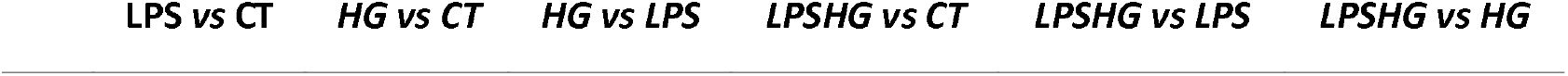
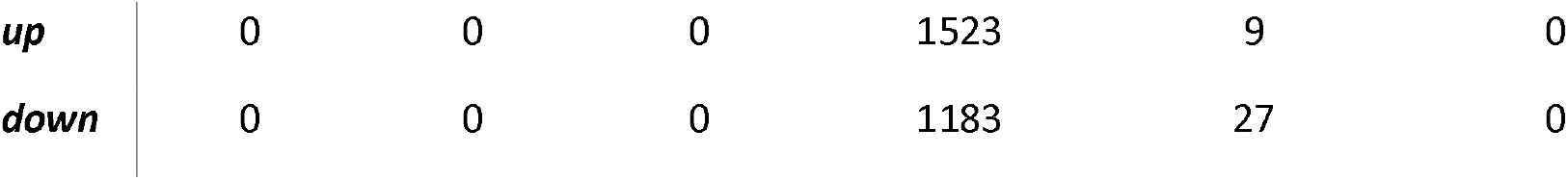
Differential Gene Expression (DGE) across the different experimental groups. The table illustrates the number of significantly upregulated and downregulated genes for each pair comparison. Groups include Control (CT), Hypergravity (HG), LPS-treated control (LPS) and LPS-treated larvae under hypergravity (LPSHG)

| LPS vs CT | HG vs CT | HG vs LPS | LPSHG vs CT | LPSHG vs LPS | LPSHG vs HG |
| --- | --- | --- | --- | --- | --- |

|  |  |  |  |  |  |  |
| --- | --- | --- | --- | --- | --- | --- |
| *up* | 0 | 0 | 0 | 1523 | 9 | 0 |
| *down* | 0 | 0 | 0 | 1183 | 27 | 0 |

No significant transcriptomic changes were detected in either the LPS or HG groups compared with CT, as all comparisons yielded zero DEGs. Likewise, no DEGs were identified between LPS and HG, indicating that neither LPS stimulation nor 3*g* hypergravity exposure alone induced detectable transcriptional alterations under the experimental conditions tested. Zero DEGs were detected in the LPSHG vs HG comparison as well.

In contrast, the combined hypergravity and LPS treatment (LPSHG) induced substantial transcriptomic changes relative to the control group. The LPSHG vs CT comparison identified a total of 2,706 DEGs (**Dataset_1**), including 1,523 upregulated and 1,183 downregulated genes. However, when LPSHG was compared with the LPS group, only 36 DEGs were detected (**Dataset_2**), of which 9 were upregulated and 27 were downregulated.

Hierarchical clustering heatmaps were generated for the two comparisons showing differential gene expression (LPSHG vs LPS and LPSHG vs CT) to visualize overall transcriptional patterns. **Figure 2a** highlights a marked transcriptomic divergence between the CT and LPSHG groups, with a total of 2,706 DEGs identified. The clustering analysis revealed a clear separation between both groups, consistent with the extensive transcriptional changes observed in the differential expression analysis. In addition, heatmap in **Figure 2b** shows LPS and LPSHG comparison. Although only 36 DEGs were identified, samples from the two groups formed distinct clusters, indicating a consistent transcriptional signature associated with the combined treatment.

**Figure 2.**
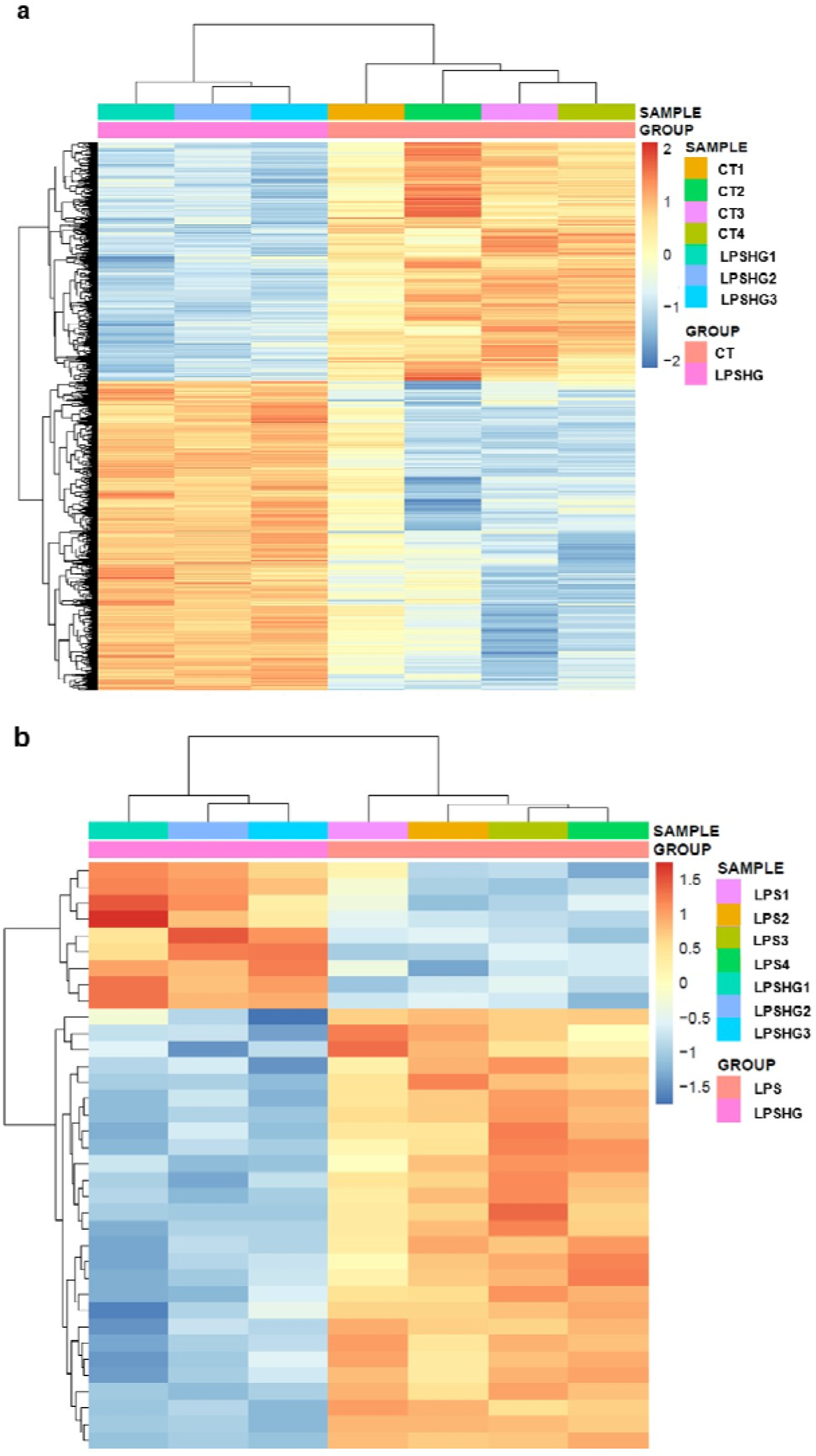
Global gene expression patterns and hierarchical clustering of differentially expressed genes (DEGs). **(a)** Heatmap illustrates the massive transcriptomic shift between the unmanipulated standard Control (CT) and the LPSHG group, encompassing 2,706 DEGs. The stark contrast in color gradients between the two clades highlights a profound, systemic physiological response. In both panels, rows represent individual genes and columns represent biological replicates. The color scale indicates normalized relative expression levels, with warmer colors (red/orange) representing higher expression and cooler colors (blue) representing lower expression. **(b)** Heatmap comparing the LPS to the dual-stress group (LPSHG), visualizing the expression profiles of the 36 significant DEGs. The distinct clustering of samples by treatment group indicates a targeted molecular response to the immunostimulation under hypergravity.

### Functional enrichment analysis and targeted selection of genes

Functional enrichment analysis of the DEGs revealed distinct biological responses. The LPSHG vs CT comparison identified 176 enriched Gene Ontology (GO) terms (**Dataset_3**), of which the top 50 are shown in **Figure 3a**. The top 10 enriched categories were the following and were predominantly associated with immunity and development: *Developmental process, Cell periphery, Anatomical structure development, Multicellular organism development, Multicellular organismal process, Protein binding, Plasma membrane, Anatomical structure morphogenesis, System development*, and *Cell differentiation*.

**Figure 3.**
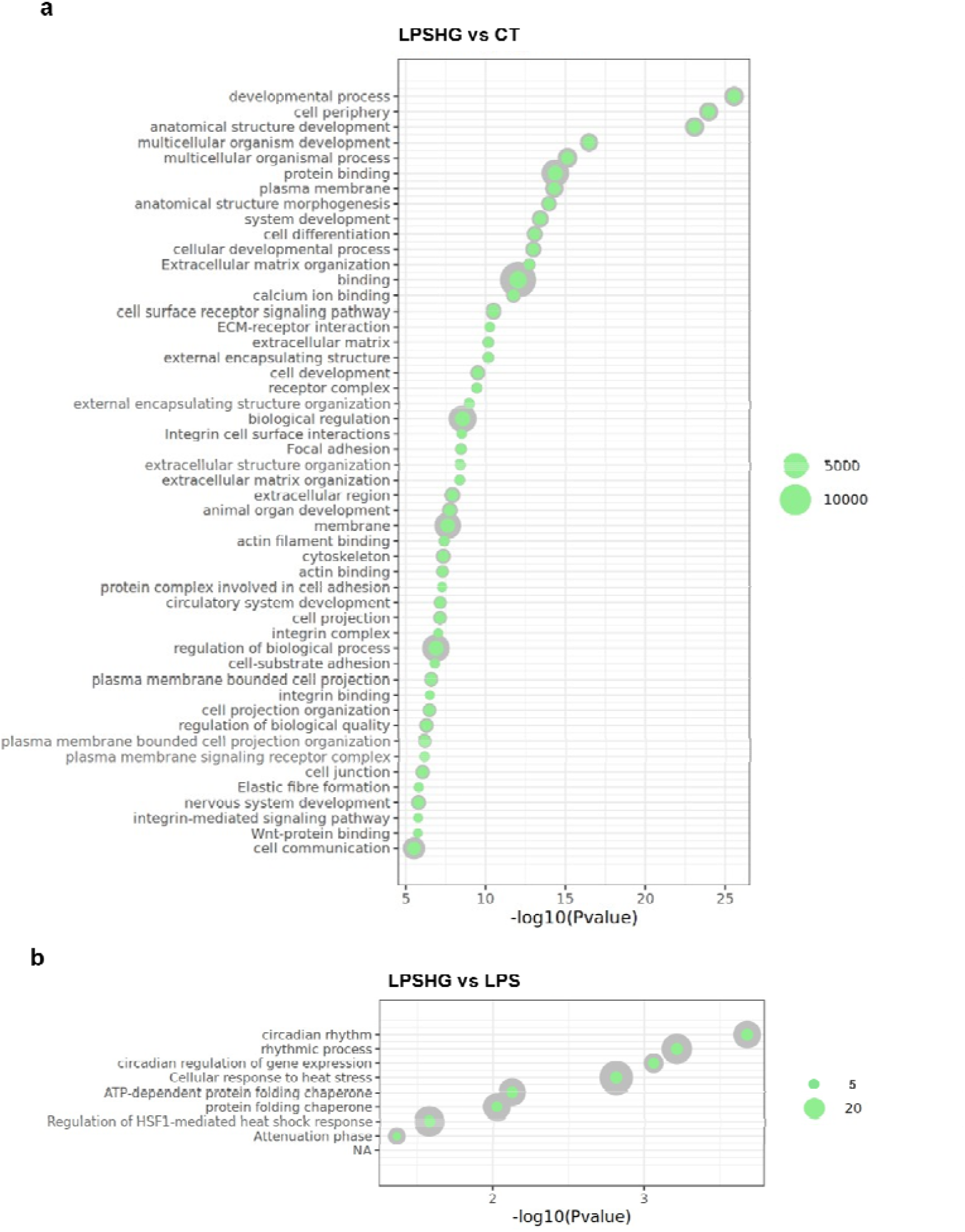
Gene Ontology (GO) enrichment analysis of differentially expressed genes. **(a)** Top 50 enriched biological processes in the LPSHG vs CT comparison, showing a broad impact on developmental, structural, and immune pathways. The x-axis represents the log2FoldChange. In each bubble, the grey outer circle represents the total number of genes associated with the pathway, while the green inner circle represents the number of those genes that were significantly differentially expressed. **(b)** Top 10 enriched biological processes in the LPSHG vs LPS comparison, highlighting the prevalence of circadian and rhythmic regulation terms.

In contrast, the LPSHG vs LPS comparison yielded a more restricted set of 8 GO terms (**Dataset_4**). The top three most significant categories were *Circadian rhythm, Rhythmic process*, and *Circadian regulation of gene expression* (**Figure 3b**).

To identify the core drivers within the LPSHG vs CT comparison, a refined statistical filter was applied (|Log_2FC| > 2, adj. p value < 0.03, **Dataset_5**), and a selection of genes related to immunity and development overlapping with the Top10 GO categories was performed.

This analysis highlighted a divergent immune response, characterized by the upregulation of genes such as *mfap4, tnfrsf1b, ptx3, alox5a, ripk1l*, and *il13r2*, alongside the downregulation of *hsp90aa1*.*2, socs1a, gpnmb*, and *cyp2r1*. Simultaneously, key developmental regulators, including *ompb, tuba8l5, mxra5, gdf5, umodl1, wnt7a, wnt9a, mia, wnt5b*, and *pxdn*, howed significant upregulation (**Figure 4a, Supplementary table S1**).

**Figure 4.**
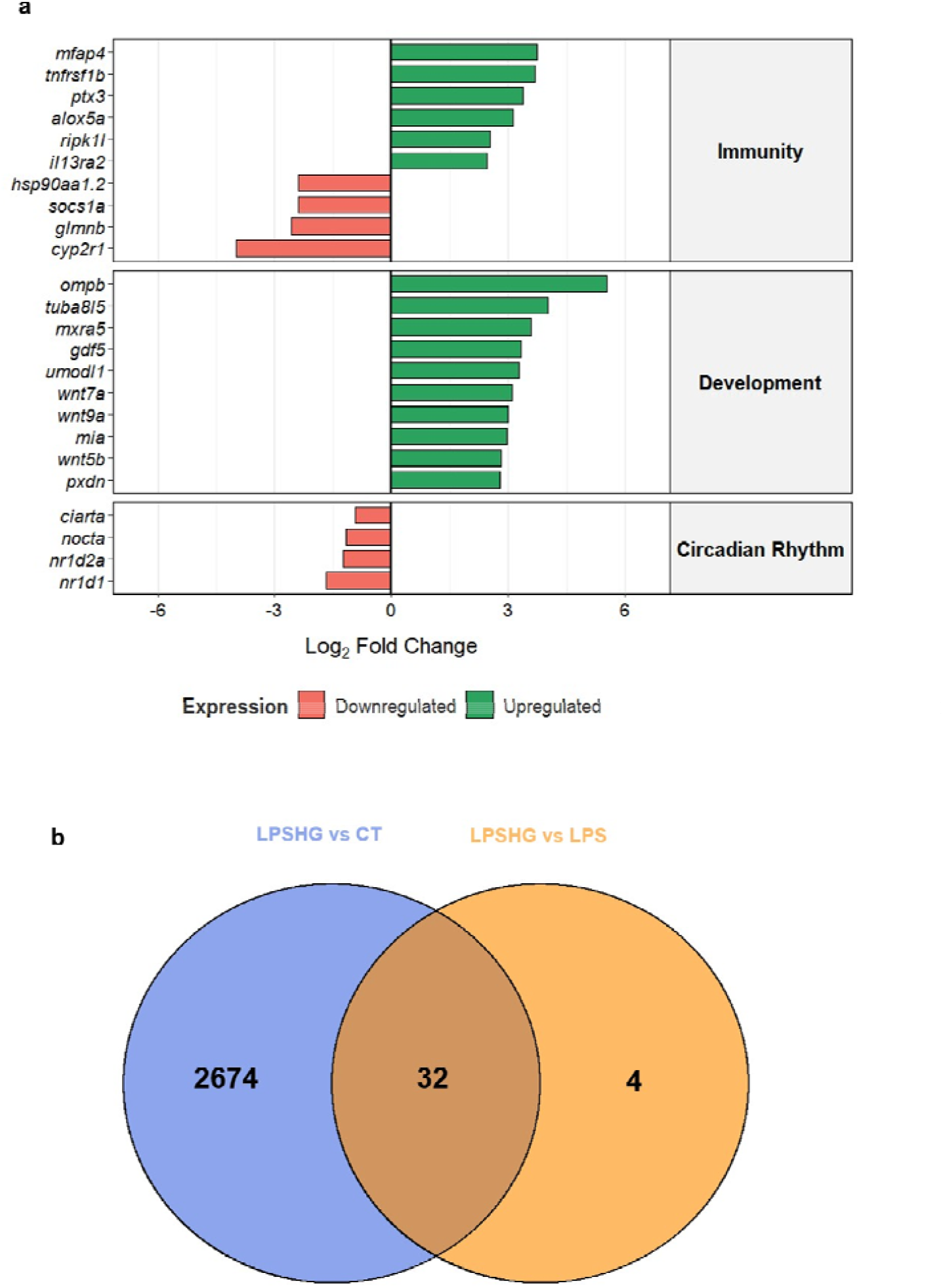
Transcriptomic fingerprint of the combined hypergravity and immunological challenge. **(a)** The horizontal bar chart summarizes the expression profiles of 24 key molecular drivers manually curated from enriched Gene Ontology (GO) categories. Immunity and development genes were selected from the overlap of the filtered LPSHG vs CT dataset (|log2FC| > 2, adj. p-value<0.03) with the Top 10-ranked GO terms of the LPSHG vs CT comparison. Circadian Rhythm genes represent the primary signature identified in the Top3 GO terms of the LPSHG vs LPS comparison. Bars represent log2 Fold Changes, with green and red indicating up- and downregulation, respectively. **(b)** Venn diagram illustrating the overlap of differentially expressed genes (DEGs) between the LPSHG vs CT (blue) and LPSHG vs LPS (orange) comparisons. The intersection reveals a robust core set of 32 genes that are differentially expressed under the LPSHG condition.

For the LPSHG vs LPS comparison, focusing on the top three circadian-related GO categories revealed a distinct subset of strongly downregulated genes, specifically *ciarta, nocta, nr1d2a*, and *nr1d1* (**Figure 4a, Supplementary table S1**). These findings indicated that the addition of the immune challenge to an immune-stimulated condition disrupts the biological clock.

Furthermore, a Venn diagram analysis was performed to determine the overlap of DEGs between the LPSHG vs CT and the LPSHG vs LPS comparisons. The intersection identified 32 shared DEGs, which represent the core mechanism of the combined hypergravity and immune challenge (LPSHG), as they are robustly altered regardless of the reference baseline (**Figure 4a, Supplementary table S2**). The function of the majority of these 32 shared genes was mostly related to immunity, but also some were related to development and circadian rhythms explaining the same molecular mechanisms driving the dual stress response.

On the other hand, the four unique genes to the LPSHG vs LPS comparison (i.e., *hspa8b, nocta, nr1d2a*, and *nupr1b*) revealed a distinct behavioral profile driven by LPS.

### Validation of RNA-seq transcriptomic profiles via qPCR

To evaluate the technical and biological reliability of the RNA-seq data, qPCR validation was performed on a selected panel of 11 candidate genes. Correlation analysis revealed a robust and highly significant positive relationship between the log2 Fold Changes obtained from the RNA-seq pipeline and the qPCR assays (**Figure 5**). The overall correlation across the entire dataset was strong (R^2^= 0.60, p = 0.00018), with consistent results observed in both experimental comparisons: LPSHG vs CT (R^2^ = 0.65, p = 0.016) and LPSHG vs LPS (R^2^ = 0.55, p = 0.014). This high degree of correlation between the two methodologies validates the transcriptomic shifts, confirming the robustness of our findings.

**Figure 5.**
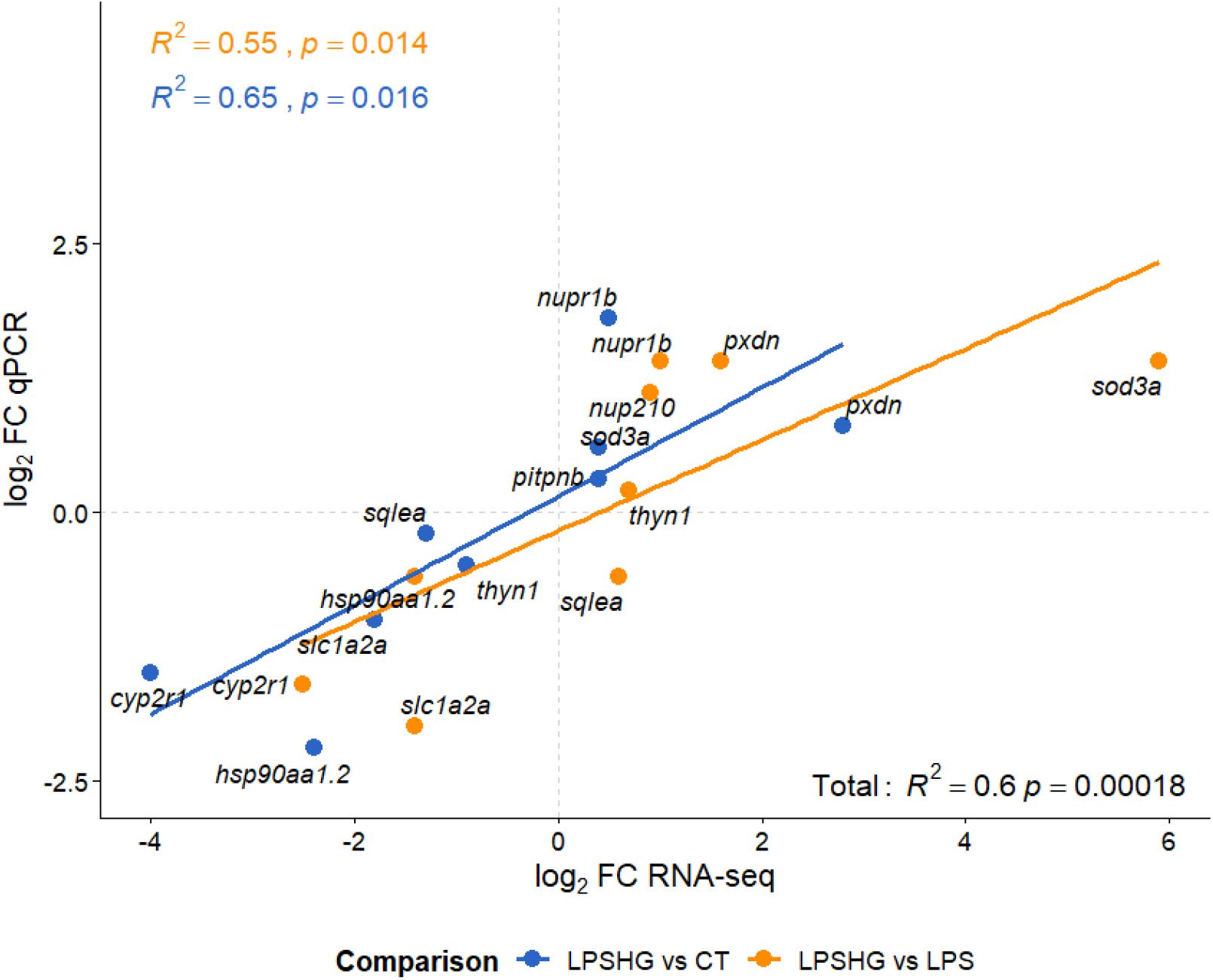
Validation of RNA-seq transcriptomic profiles via quantitative PCR (qPCR). The scatter plot illustrates the correlation between log2 Fold Changes measured by RNA-seq (x-axis) and qPCR (y-axis) for 11 selected genes. Regression lines are shown for the LPSHG vs CT (blue) and LPSHG vs LPS (orange) comparisons. A significant positive correlation was observed across the dataset (R^2^ = 0.60, p < 0.001), confirming the reliability of the high-throughput sequencing results.

## Discussion

The present study provides for the first time a comprehensive transcriptomic analysis of the synergistic effects of gravity and immune system alterations in fish. While ground-based hypergravity models are valuable for deciphering how gravitational changes influence physiological mechanisms, the interaction between gravitational stress and innate immunity at the transcriptomic level has never been previously explored. Here, the establishment of an optimal immunological challenge was set down in the 1–10□µg/ml range, a dose that strikes a biological balance: triggering a robust, measurable inflammatory response while keeping over 70% larval viability. This non-lethal stimulation is key for space biology and aquaculture models, allowing to study long-term physiological adaptation and underlying molecular mechanisms.

One of the most striking findings of the current study is the transcriptomic stability of zebrafish larvae exposed to single stressors. Neither the 3*g* hypergravity condition (HG) nor the LPS stimulation independently resulted in detectable transcriptional alterations compared to the standard control. This lack of transcriptomic divergence is in accordance with gene expression studies previously reported in the same animal models, in which no significance was found in some key genes related to epigenetics ^15^. In contrast, a previous study ^14^ observed that exposing 5 days zebrafish larvae to simulated microgravity induced significant baseline transcriptomic shifts, particularly in stress and musculoskeletal pathways. The fact that neither the 3*g* hypergravity condition nor the LPS stimulation resulted in 0 DEGs suggestes that zebrafish exhibits remarkable transcriptomic resilience to hypergravity and LPS alone and is an advantageous model for future space-biological research. By understanding how these organisms maintain transcriptomic stability under 3*g*, we can learn valuable molecular strategies for gravity adaptation for humans.

However, the combined exposure to LPS in the minimal tested dose and hypergravity (LPSHG) resulted in a marked transcriptomic response relative to the standard control group (CT), with a total of 2,706 differentially expressed genes (1,523 upregulated and 1,183 downregulated). This extensive transcriptional remodeling contrasts sharply with the absence of significant changes observed when either HG or LPS alone was compared with the control group. Thus, the results were able to characterize the earliest transcriptomic changes associated with a mild immune response.

Interestingly, the comparison between LPSHG and the LPS group identified only 36 DEGs. This indicated that even though the dual-stress condition compared to the control caused a high systemic shock, the addition of the hypergravity to the immune challenge elicited only a highly targeted response, suggesting that the immune challenge is the primary determinant of the transcriptomic changes.

In addition, by intersecting the LPSHG vs CT and LPSHG vs LPS comparisons with the Venn diagram, we isolated a core mechanism of 32 shared DEGs. These genes may have represented specific pathways involved in the interaction between gravitational stress and immune activation. Overall, the two comparisons seemed to be explanatory of the common molecular mechanism driving the dual-stress condition. Notably, four genes (*hspa8b, nocta, nr1d2a, nupr1b*) were uniquely expressed in the LPSHG vs LPS comparison, suggesting that they acted as highly specific responders to hypergravity stress during infection.

Unexpectedly, no DEGs were detected in the comparison between LPSHG and HG. While this pattern may indicate a complex interaction between hypergravity and immune stimulation, the absence of detectable differences should be interpreted with caution, as it appears difficult to reconcile with the large number of DEGs observed in the LPSHG vs CT comparison. Further analyses will therefore be required to determine whether this result reflects a biological phenomenon or methodological limitations associated with the statistical analysis. Nevertheless, the substantial transcriptomic response observed in the LPSHG group prompted us to investigate the specific molecular pathways underlying this dual-stress condition.

Functional characterization and GO enrichment analyses in the two comparisons revealed three major interconnected biological modules: immune regulation, developmental remodeling and circadian pathways. Collectively, these pathways provide insights into the molecular mechanisms through which altered gravity may influence physiological adaptation during immune challenge.

The interaction between altered gravity and immune function is of particular relevance for long-duration space missions. Extensive clinical and physiological monitoring of astronauts demonstrated that exposure to the space environment induces persistent immune dysregulation and increases susceptibility to infections ^35^. Our zebrafish model captured potential molecular mechanisms underlying this vulnerability. Unlike conventional LPS responses observed under standard Earth gravity, which typically induce broad activation of pathogen recognition pathways, pathogen clearance mechanisms and inflammasome-associated responses ^36^, the dual-stress condition produced a more selective pattern of immune remodeling.

Several genes involved and immune regulation, were selected from the GO term categories of the LPSHG vs CT comparison. The most upregulated gene was *mfap4*, which encodes a protein active in the extracellular space and as demonstrated in a previous study, it serves as a very specific promoter for macrophages in zebrafish ^37^.

The differential expression of *mfap4* in our model indicates that the combined stressors significantly alter macrophage dynamics and the extracellular matrix. Moreover, we observed a robust upregulation of *tnfrsf1b*a primary driver of pro-inflammatory cytokine signaling. Recent studies have demonstrated that *tnfrsf1b* played a critical protective role in zebrafish during bacterial infections by mediating host-pathogen interactions and regulating macrophage survival ^38^. Another interesting, upregulated gene is *ptx3*, which is known to act as a crucial link between innate immunity and tissue remodelling in zebrafish, and is a biomarker of acute bacterial infection in fish and humans ^39,40^.

In teleost fish, including zebrafish, cytokines are primary orchestrators of the immune response, initiating inflammation following exposure to pathogens or pathogen-associated molecular patterns like LPS ^41^. However, the transcriptomic profile of this study revealed a profound disruption within cytokine signaling. Interestingly, we observed a significant downregulation of the suppressor *sosc1a*, which functions as a negative regulator for cytokine pathways ^42^. Moreover, we observed an upregulation of *il13ra2*, a receptor involved in the modulation of the cytokine interleukin-13 and inflammatory responses ^43^. These modifications related to cytokines may suggest fundamental changes in the immune activity under these studied conditions.

Particularly noteworthy was the strong downregulation of *cyp2r1*, a key enzyme involved in vitamin D metabolism. Vitamin D signaling is essential for maintaining immune homeostasis and limiting excessive inflammatory responses in teleost fish ^44^. Consequently, suppression of *cyp2r1* may reduce the ability of larvae to regulate inflammation appropriately under altered gravity conditions. Together, these findings indicate that hypergravity does not simply enhance immune activity but instead reshapes the architecture of the immune response during immune challenge.

Beyond immune dysregulation, the functional analysis of the LPSHG vs CT comparison revealed a profound upregulation of genes associated with extensive developmental, cytoskeletal and structural remodeling. Previous studies have demonstrated that hypergravity exposure affects cartilage formation, craniofacial morphology and skeletal development in fish ^9,16,17^. Our transcriptomic observations over the 5-day experimental period provide the molecular background for these reported phenotypes. Across the most prominent developmental regulators were multiple members of the wnt signaling pathway (*wnt5b, wnt7a, wnt9a*) which play central roles in tissue patterning, morphogenesis and skeletal development ^45^. The simultaneous upregulation of *gdf5* -a gene crucial for development and patterning of the endoskeleton, including joints and fin radials-further supports these findings Notably, based on zebrafish database (ZFIN) annotations, *gdf5, wnt7a*, and *wnt9a* were also predicted to enable cytokine activity. This dual functionality suggested that their robust upregulation was not exclusively structural; rather, it actively bridges tissue remodeling with the ongoing inflammatory response. Together, their robust upregulation suggested an active initiation of tissue repair and remodeling triggered by dual stress, indicating that hypergravity exposure during immune challenge significantly interferes with normal developmental programs and tissue maintenance. Furthermore, beyond skeletal changes, the larvae seemed to be actively reinforcing their cellular architecture. A significant upregulation of extracellular matrix (ECM) and cytoskeletal genes was observed, such as *pxdn* and *mxra5*. At the molecular level, *pxdn* is required to stabilize collagen networks in the ECM by forming structural cross-links ^46^. Similarly, *mxra5* was involved in active-matrix remodeling during tissue injury and inflammation and was considered an immune-biomarker. Overall, this coordinated genetic response implied that hypergravity exposure during an immune challenge forced the organism to divert significant biological resources toward reinforcing tissue integrity, fundamentally altering standard developmental programs.

The final component of the analysis was the circadian rhythm related genes, which were identified in the LPSHG vs LPS comparison: *ciarta, nocta, nr1d2a, nr1d1*. The disruption of circadian regulation is particularly relevant because circadian rhythms are tightly interconnected with physiological adaptation and immune function. Altered gravity has previously been associated with disturbances in biological rhythms and behavioural responses in aquatic organisms ^47^. Similarly, exposure to altered gravity has been shown to impair vestibular function and induce physiological stress in fish species ^9^. Since circadian pathways are deeply intertwined with immune regulation, the observed suppression of *nr1d1* and *ciarta* genes suggested that hypergravity may indirectly influence immune competence through disruption of temporal homeostasis. In zebrafish, suppression of *nr1d1* was linked to the establishment of a pro-inflammatory microenvironment ^48^, providing a potential mechanistic explanation for the immune dysregulation observed under dual-stress conditions. Together, these findings suggested that the dual condition affected not only immune pathways directly but also key regulatory systems that coordinate physiological adaptation to environmental stress. Even more strikingly, the Venn diagram analysis allowed us to isolate four genes exclusively altered in the LPSHG vs LPS comparison (*nocta, nr1d2a, hspa8b*, and *nupr1b*). These genes exhibited a transcriptional profile driven entirely by the addition of gravitational stress to the infected larvae. The downregulation of *nocta* and *nr1d2a* showed the profound impact of hypergravity on circadian rhythm and further homeostasis.

Taken together, our results indicated that hypergravity alone was largely tolerated at the transcriptomic level, whereas its combination with immune stimulation triggered a coordinated reorganization of immune signaling development and circadian regulation. The genes identified and studied here consisted of potential biomarkers and therefore represented a promising molecular framework for understanding how vertebrate organisms adapt to the combined challenges of altered gravity and infection, two conditions that are highly relevant for future long-duration space missions.

## Conclusions

In conclusion, this study successfully established a novel dual-stress model combining 3*g* hypergravity and an LPS-induced immune challenge over a 5-day observation period in zebrafish larvae. While this organism showed exceptional transcriptomic resilience to either hypergravity or LPS independently, the combination of hypergravity and immune stimulation triggered extensive systemic remodeling. When the larvae faced both stressors simultaneously, three biological systems were profoundly affected: immunity, development, and circadian rhythm. First, the dual-stress condition disrupted normal immune tolerance and reshaped cytokine signaling. This was driven by the suppression of protective genes (*socs1a, cyp2r1*) alongside the upregulation of receptors (like *il13ra2*), which points to a fundamental alteration in inflammatory modulation. Second, this combined gravitational and immune challenge caused structural and tissue remodeling. This developmental shift was highlighted by the upregulation of skeletal master regulators (*wnt5b, wnt7a and gdf5*) and extracellular matrix stabilizers (*pxdn, mxra5*). Notably, because genes like *gdf5* and *wnt7a* were also predicted to enable cytokine activity, their activation actively bridges structural tissue repair with the ongoing inflammatory response. Finally, the synergistic stress affected the internal biological clock by downregulating core circadian repressors (*nr1d1, ciarta*), ultimately exposing the organism to systemic stress. Overall, these results suggested that the combination of immune challenge and hypergravity compromised immune tolerance, forced developmental remodeling, and disrupted circadian rhythm. Understanding these specific molecular vulnerabilities in space-related environments in model animals will be essential for protecting astronauts’ health and ensuring the success and sustainability of future space aquaculture.

## Author contributors

**MM:** Writing – original draft, Visualization, Formal Analysis, Investigation, Data curation. **EC:** Software, Data curation, Formal Analysis, Writing – review and editing. **AE:** Software, Data curation, Formal Analysis, Writing – review and editing. **LR:** Conceptualization, Supervision, Resources, Project administration, Funding acquisition, Writing – original draft.

## Acknowledgements

This study was funded through Agencia Espacial Española (AEE) and implemented by the European Space Agency (ESA, ESA4000148975) PRODEX Programme, SPACEGENFISH, to LR and with funding from the Spanish government through the ‘. We thank Marcela Salazar, Sílvia Joly and Dario Iruela Ortiz for helping us with the experiment and laboratory analyses. Institutional support to CNAG was provided by the Spanish Ministry of Science and Innovation through the Instituto de Salud Carlos III, and by the Generalitat de Catalunya through the Departament de Salut and the Departament de Recerca i Universitats.

## Contents of Datasets

Dataset_1: DEGs of LPSHG vs CT comparison

Dataset_2: DEGs of LPSHG vs LPS comparison

Dataset_3: GO terms of LPSHG vs CT comparison

Dataset_4: GO terms of LPSHG vs LPS comparison

Dataset_5: Filtered DEGs LPSHG vs CT comparison, to overlap with the functional enrichment analysis.

## Bibliography

1. Volkmann, D. & Baluška, F. Gravity: One of the driving forces for evolution. Protoplasma 229, 143–148 (2006).

2. Wade, C. E. Responses across the Gravity Continuum: Hypergravity to Microgravity. Advances in Space Biology and Medicine vol. 10 225–245 Preprint at 10.1016/S1569-2574(05)10009-4 (2005).

3. Frett, T., Petrat, G., Jack, J. J., Hemmersbach, R. & Anken, R. Hypergravity Facilities in the ESA Ground-Based Facility Program – Current Research Activities and Future Tasks. Microgravity Sci. Technol. 28, 205–214 (2016).

4. Hariom, S. K. et al. Animal physiology across the gravity continuum. Acta Astronautica vol. 178 522–535 Preprint at 10.1016/j.actaastro.2020.09.044 (2021).

5. Morita, H. et al. Feasibility of a short-arm centrifuge for mouse hypergravity experiments. PLoS One 10, (2015).

6. Marcu, O. et al. Innate immune responses of Drosophila melanogaster are altered by spaceflight. PLoS One 6, (2011).

7. Frippiat, J. P. Contribution of the urodele amphibian Pleurodeles waltl to the analysis of spaceflight-associated immune system deregulation. Molecular Immunology vol. 56 434–441 Preprint at 10.1016/j.molimm.2013.06.011 (2013).

8. Ohnishi, T. et al. Accumulation of stress protein 72 (HSP72) in muscle and spleen of goldfish taken into space. Ah. Space Res vol. 21 (1998).

9. Tanaka, K., Nishimura, N. & Kawai, Y. Adaptation to microgravity, deconditioning, and countermeasures. Journal of Physiological Sciences vol. 67 271–281 Preprint at 10.1007/s12576-016-0514-8 (2017).

10. Sonnenfeld, G. & New Collective Author. Use of animal models for space flight physiology studies, with special focus on the immune system. Gravit. Space Biol. Bull. 18, 31–5 (2005).

11. Spence, R., Gerlach, G., Lawrence, C. & Smith, C. The behaviour and ecology of the zebrafish, Danio rerio. Biological Reviews vol. 83 13–34 Preprint at 10.1111/j.1469-185X.2007.00030.x (2008).

12. Ribas, L. & Piferrer, F. The zebrafish (Danio rerio) as a model organism, with emphasis on applications for finfish aquaculture research. Reviews in Aquaculture vol. 6 209–240 Preprint at 10.1111/raq.12041 (2014).

13. Piferrer, F. & Ribas, L. The use of the zebrafish as a model in fish aquaculture research. in Fish Physiology vol. 38 273–313 (Elsevier Inc., 2020).

14. Aceto, J. et al. Zebrafish bone and general physiology are differently affected by hormones or changes in gravity. PLoS One 10, (2015).

15. Salazar, M., Joly, S., Anglada-Escudé, G. & Ribas, L. Epigenetic and physiological alterations in zebrafish subjected to hypergravity. PLoS One 19, (2024).

16. Moorman, S. J., Cordova, R. & Davies, S. A. A critical period for functional vestibular development in zebrafish. Developmental Dynamics 223, 285–291 (2002).

17. Lawrence, E. A. et al. Exposure to hypergravity during zebrafish development alters cartilage material properties and strain distribution. Bone Joint Res. 10, 137–148 (2021).

18. Garrett-Bakelman, F. E. et al. The NASA twins study: A multidimensional analysis of a year-long human spaceflight. Science (1979). 364, (2019).

19. Grigoryan, E. N. Impact of Microgravity and Other Spaceflight Factors on Retina of Vertebrates and Humans In Vivo and In Vitro. Life vol. 13 Preprint at 10.3390/life13061263 (2023).

20. Murata, Y. et al. Histological and transcriptomic analysis of adult Japanese Medaka sampled onboard the international space station. PLoS One 10, (2015).

21. Rykova, M. P., Antropova, E. N., Larina, I. M. & Morukov, B. V. Humoral and cellular immunity in cosmonauts after the ISS missions. Acta Astronaut. 63, 697–705 (2008).

22. Dobney, W. et al. Evaluation of deep space exploration risks and mitigations against radiation and microgravity. Frontiers in Nuclear Medicine vol. 3 Preprint at 10.3389/fnume.2023.1225034 (2023).

23. Hurem, S. et al. Dose-dependent effects of gamma radiation on the early zebrafish development and gene expression. PLoS One 12, (2017).

24. Campisi, M., Cannella, L. & Pavanello, S. Cosmic chronometers: Is spaceflight a catalyst for biological ageing? Ageing Research Reviews vol. 95 Preprint at 10.1016/j.arr.2024.102227 (2024).

25. Munford, R. S. Sensing gram-negative bacterial lipopolysaccharides: A human disease determinant? Infection and Immunity vol. 76 454–465 Preprint at 10.1128/IAI.00939-07 (2008).

26. Novoa, B., Bowman, T. V., Zon, L. & Figueras, A. LPS response and tolerance in the zebrafish (Danio rerio). Fish Shellfish Immunol. 26, 326–331 (2009).

27. OECD/OCDE 234. OECD guideline for the testing of chemicals. Fish Sexual Development Test. Introduction. (2011).

28. Dobin, A. et al. STAR: Ultrafast universal RNA-seq aligner. Bioinformatics 29, 15–21 (2013).

29. Li, B. & Dewey, C. N. RSEM: Accurate transcript quantification from RNA-Seq data with or without a reference genome. BMC Bioinformatics 12, (2011).

30. Ritchie, M. E. et al. Limma powers differential expression analyses for RNA-sequencing and microarray studies. Nucleic Acids Res. 43, e47 (2015).

31. Law, C. W., Chen, Y., Shi, W. & Smyth, G. K. Voom: Precision Weights Unlock Linear Model Analysis Tools for RNA-Seq Read Counts. Genome Biology vol. 15 http://genomebiology.com/2014/15/2/R29 (2014).

32. Kolberg, L. et al. G:Profiler-interoperable web service for functional enrichment analysis and gene identifier mapping (2023 update). Nucleic Acids Res. 51, W207–W212 (2023).

33. Korotkevich, G. et al. Fast gene set enrichment analysis. Preprint at 10.1101/060012 (2016).

34. Oliveros, J. C. Venny. An interactive tool for comparing lists with Venn’s diagrams. (2015).

35. Crucian, B. E. et al. Immune system dysregulation during spaceflight: Potential countermeasures for deep space exploration missions. Frontiers in Immunology vol. 9 Preprint at 10.3389/fimmu.2018.01437 (2018).

36. Watzke, J., Schirmer, K. & Scholz, S. Bacterial lipopolysaccharides induce genes involved in the innate immune response in embryos of the zebrafish (Danio rerio). Fish Shellfish Immunol. 23, 901–905 (2007).

37. Walton, E. M., Cronan, M. R., Beerman, R. W. & Tobin, D. M. The macrophage-specific promoter mfap4 allows live, long-term analysis of macrophage behavior during mycobacterial infection in zebrafish. PLoS One 10, (2015).

38. Jing, Y. et al. Tnfrsf1b Protects Zebrafish Against Klebsiella pneumoniae Infection. Infect. Drug Resist. 18, 5473–5488 (2025).

39. Doni, A. et al. The long pentraxin PTX3 as a link between innate immunity, tissue remodeling, and cancer. Frontiers in Immunology vol. 10 Preprint at 10.3389/fimmu.2019.00712 (2019).

40. Bello-Perez, M. et al. Modulation of the tissue expression pattern of zebrafish crp-like molecules suggests a relevant antiviral role in fish skin. Biology (Basel). 10, 1–13 (2021).

41. Zou, J. & Secombes, C. J. The function of fish cytokines. Biology vol. 5 Preprint at 10.3390/biology5020023 (2016).

42. Zhang, Y. et al. The Roles of SOCS in Teleost Immunity: Cytokine Regulation and Host Defense. Reviews in Aquaculture vol. 18 Preprint at 10.1111/raq.70161 (2026).

43. Bottiglione, F. et al. Zebrafish IL-4–like Cytokines and IL-10 Suppress Inflammation but Only IL-10 Is Essential for Gill Homeostasis. The Journal of Immunology 205, 994–1008 (2020).

44. Liao, X. et al. Vitamin D influences gut microbiota and acetate production in zebrafish (Danio rerio) to promote intestinal immunity against invading pathogens. Gut Microbes 15, (2023).

45. Polevoy, H. et al. New roles for Wnt and BMP signaling in neural anteroposterior patterning. The EMBO Reports 20, EMBR201845842 (2019).

46. Fidler, A. L. et al. A unique covalent bond in basement membrane is a primordial innovation for tissue evolution. Proc. Natl. Acad. Sci. U. S. A. 111, 331–336 (2014).

47. Gibbs, J. E. et al. The nuclear receptor REV-ERBα mediates circadian regulation of innate immunity through selective regulation of inflammatory cytokines. Proc. Natl. Acad. Sci. U. S. A. 109, 582–587 (2012).

48. Wu, M. et al. Role of NR1D1 in Bisphenol A-Induced Anxiety-like Behavior and Inflammation in Zebrafish Larvae. Toxics 13, (2025).

