## Supplementary tables for "Synergistic effects of hypergravity and immune challenge on the zebrafish larval transcriptome": Supplem tables_Mitsopoulou_Submited.docx

**Table S1:** Detailed list of the 24 selected genes associated with immunity, development, and circadian rhythms. Values reflect log2 Fold Changes (log2FC) and adjusted p-values derived from the LPSHG vs CT and LPSHG vs LPS comparisons

| **Gene ID** | **Abbreviation** | **Gene name** | **log2FC** | **adj. p-value** | **Description (ZFIN)** |
| --- | --- | --- | --- | --- | --- |
| ENSDARG00000038683 | *mfap4* | Microfibril associated protein 4 | 3.740 | 0.019 | Elastic fiber assembly; extracellular matrix. Expressed in endothelium, hematopoietic system, macrophages, nervous system, and trunk. Human ortholog: MFAP4. |
| ENSDARG00000070165 | *tnfrsf1b* | Tumor necrosis factor receptor superfamily, member 1B | 3.693 | 0.027 | Signaling receptor (plasma membrane); hematopoietic differentiation, PNS axon ensheathment, viral response. Expression: head, hematopoietic system, osteoclasts, scales. Disease model: psoriasis. Human ortholog: TNFRSF1B (Parkinsonism; acne; bone disease; glomerulonephritis; and lung disease). |
| ENSDARG00000068275 | *ptx3* | Pentraxin 3 | 3.376 | 0.023 | C1q complex binding and innate immune response (extracellular space). Expressed in epidermis, fins, hypochord, and nervous system. Human ortholog: PTX3 (cystic fibrosis, invasive aspergillosis). |
| ENSDARG00000057273 | *alox5a* | Arachidonate 5-lipoxygenase a | 3.126 | 0.023 | Arachidonate 5-lipoxygenase/monooxygenase and leukotriene-C4 synthase activity (cytoplasm/nuclear envelope). Inflammatory response to wounding. Expressed in pronephric duct. Human ortholog: ALOX5 (artery disease, asthma, pancreatic cancer, pulmonary tuberculosis). |
| ENSDARG00000006677 | *ripk1l* | Receptor (TNFRSF)-interacting serine-threonine kinase 1, like | 2.536 | 0.024 | JUN kinase activity / signal transduction (cytoplasm). Expression: head, heart, integument, liver, pleuroperitoneum. Human ortholog: RIPK1 (immunodeficiency 57). |
| ENSDARG00000039436 | *il13ra2* | Interleukin 13 receptor, alpha 2 | 2.460 | 0.022 | Cytokine receptor/binding activity and signaling, positive regulation of cell proliferation (plasma membrane receptor complex). Expressed in the cardiovascular system and otic vesicle. Human ortholog: IL13RA2. |
| ENSDARG00000024746 | *hsp90aa1.2* | Heat shock protein 90, alpha (cytosolic), class A member 1, tandem duplicate 2 | -2.383 | 0.018 | ATP binding/hydrolysis and unfolded protein binding (cytoplasm, myelin sheath, neuronal cell bodies). Involved in protein stabilization and cellular response to heat/cold. Expressed in immature eye, mesoderm, musculature, nervous system, and pericardial region. Human ortholog: HSP90AA1. |
| ENSDARG00000038095 | *socs1a* | Suppressor of cytokine signaling 1a | -2.402 | 0.023 | Cytokine receptor binding / STAT phosphorylation / T cell migration. Expression: brain, digestive/musculature/visual systems, pleuroperitoneum. Disease models: hyperinsulinism, steatotic liver disease. Human ortholog: SOCS1 (leukemia, multiple cancers). |
| ENSDARG00000010958 | *glmnb* | Glomulin, FKBP associated protein b | -2.583 | 0.018 | Ubiquitin-protein transferase inhibitor activity (cytoplasm). Human ortholog: GLMN (implicated in arteriovenous malformations, glomangiomas, and glomus tumors). |
| ENSDARG00000056587 | *cyp2r1* | Cytochrome P450, family 2, subfamily R, polypeptide 1 | -3.986 | 0.018 | Vitamin D3 25-hydroxylase / heme binding (ER membrane/cytoplasm); vitamin D metabolism, hematopoietic proliferation. Expression: liver, muscle, ovary, visceral fat. Disease model: vitamin D-dependent rickets 1B. Human ortholog: CYP2R1 (implicated in obesity, type 1 diabetes, brain infarction, artery disease, and vitamin D-dependent rickets type 1B). |
| ENSDARG00000032380 | *ompb* | Olfactory marker protein b | 5.531 | 0.027 | Is expressed in olfactory system and sensory neurons. Orthologous to human OMP. |
| ENSDARG00000094466 | *tuba8l5* | Tubulin, alpha 8 like 5 | 4.015 | 0.022 | GTP binding and structural constituent of the cytoskeleton (cytoplasm/ microtubules). Involved in microtubule organization and the mitotic cell cycle. |
| ENSDARG00000076309 | *mxra5* | Matrix-remodelling associated 5 | 3.591 | 0.022 | Predicted to be involved in nervous system development. Predicted to be located in plasma membrane. Orthologous to human MXRA5. |
| ENSDARG00000002760 | *gdf5* | Growth differentiation factor 5 | 3.327 | 0.023 | Cytokine activity (extracellular space). Involved in medial/pectoral fin and skeletal system morphogenesis. Expressed in perichondrium, pharynx, and skeletal system. Human ortholog: GDF5 (Parkinson's, bone diseases). |
| ENSDARG00000068923 | *umodl1* | Uromodulin-like 1 | 3.280 | 0.025 | Epithelial cell differentiation (cell surface and extracellular space). Human ortholog: UMODL1. |
| ENSDARG00000002483 | *wnt7a* | Wingless-type MMTV integration site family, member 7A | 3.104 | 0.022 | Predicted to enable Cytokine activity/frizzled binding, canonical Wnt signaling, neuron differentiation, and positive regulation of the JNK cascade (extracellular space). Expressed in brain, head, otic vesicle, pectoral fin bud, and spinal cord. Human ortholog: WNT7A (implicated in Fuhrmann syndrome and Schinzel type phocomelia). |
| ENSDARG00000063437 | *wnt9a* | Wingless-type MMTV integration site family, member 9A | 3.007 | 0.027 | Cytokine/frizzled binding, embryonic heart tube formation, hematopoietic stem cell differentiation, and skeletal system development (extracellular space). Expressed in gut, head, nervous system, pectoral fin, and pleuroperitoneal region. Human ortholog: WNT9A (implicated in pancreatic cancer). |
| ENSDARG00000052437 | *mia* | MIA SH3 domain containing | 2.967 | 0.022 | Predicted to be involved in extracellular matrix organization. Orthologous to human MIA. |
| ENSDARG00000102464 | *wnt5b* | Wingless-type MMTV integration site family, member 5b | 2.813 | 0.022 | Signaling receptor binding, non-canonical Wnt signaling, convergent extension (gastrulation), muscle/skeletal/vasculature development, and midbrain-hindbrain boundary morphogenesis (extracellular space). Expressed in central nervous system, cranial cartilage, fins, germ ring, and mesoderm. Human ortholog: WNT5B. |
| ENSDARG00000098733 | *pxdn* | Peroxidasin | 2.799 | 0.018 | Predicted to enable heme binding activity; metal ion binding activity; and peroxidase activity. Acts upstream of or within basement membrane assembly. In endoplasmic reticulum and extracellular region. |
| ENSDARG00000058094 | *ciarta* | Circadian associated repressor of transcription a | -0.930 | 0.039 | RNA polymerase II sequence-specific DNA binding, circadian gene regulation, and negative regulation of transcription (nucleus). Expressed in pronephric duct. Human ortholog: CIART |
| ENSDARG00000077726 | *nocta* | Nocturnin a | -1.170 | 0.038 | 3'-5'-RNA exonuclease activity and nucleobase-containing compound metabolism (cytoplasm). Human ortholog: NOCT. |
| ENSDARG00000003820 | *nr1d2a* | Nuclear receptor subfamily 1, group D, member 2a | -1.252 | 0.038 | Nuclear receptor activity, RNA polymerase II sequence-specific DNA binding, circadian gene regulation, and regulation of transcription (cytoplasm/nucleus). Expressed in the nervous system. Human ortholog: NR1D2 |
| ENSDARG00000033160 | *nr1d1* | Nuclear receptor subfamily 1, group d, member 1 | -1.665 | 0.020 | DNA-binding transcriptional repressor, RNA polymerase II sequence-specific DNA binding, autophagy, circadian gene regulation, and regulation of locomotor rhythms (chromatin). Expressed in the nervous system. Human ortholog: NR1D1 (implicated in brain small vessel disease). |

**Table S2:** List of the 32 common genes between the LPSHG vs CT and the LPSHG vs LPS comparison.

| **Gene ID** | **Gene Abbreviation** | **Gene Name** | **log2FC (LPSHG vs CT)** | **Adj. p-value (LPSHG vs CT)** | **log2FC (LPSHG vs LPS)** | **Adj. p-value (LPSHG vs LPS)** |
| --- | --- | --- | --- | --- | --- | --- |
| ENSDARG00000056587 | *cyp2r1* | Cytochrome P450, family 2, subfamily R, polypeptide 1 | -4.635 | 0.017 | -4.635 | 0.017 |
| ENSDARG00000024746 | *hsp90aa1.2* | Heat shock protein 90 alpha family class A member 1, tandem duplicate 2 | -2.383 | 0.018 | -2.834 | 0.017 |
| ENSDARG00000061602 | *ssx2ip* | Synovial sarcoma, X breakpoint 2 interacting protein | -1.321 | 0.018 | -1.403 | 0.017 |
| ENSDARG00000035861 | *-* | Uncharacterized protein | -1.349 | 0.018 | -1.437 | 0.017 |
| ENSDARG00000030758 | *guca1c* | Guanylate cyclase activator 1C | -1.748 | 0.018 | -1.888 | 0.018 |
| ENSDARG00000051970 | *-* | Uncharacterized protein | -1.051 | 0.02 | -1.247 | 0.018 |
| ENSDARG00000052138 | *slc1a2a* | Solute carrier family 1 member 2a | -1.718 | 0.018 | -1.579 | 0.018 |
| ENSDARG00000099470 | *muc5.3* | Mucin 5.3 | 2.569 | 0.012 | 2.042 | 0.018 |
| ENSDARG00000019874 | *hsph1* | Heat shock protein family H (Hsp110) member 1 | -1.332 | 0.022 | -1.735 | 0.018 |
| ENSDARG00000007241 | *mycb* | Myelocytomatosis viral oncogene homolog b | -0.766 | 0.03 | -1.297 | 0.018 |
| ENSDARG00000019274 | *rasd1* | Ras dexamethasone-induced 1 | -0.983 | 0.025 | -1.463 | 0.02 |
| ENSDARG00000033160 | *nr1d1* | Nuclear receptor subfamily 1 group D member 1 | -1.157 | 0.023 | -1.665 | 0.02 |
| ENSDARG00000098164 | *-* | Uncharacterized protein | -2.025 | 0.018 | -2.008 | 0.024 |
| ENSDARG00000058285 | *cpt1b* | Carnitine palmitoyltransferase 1B | 1.5 | 0.022 | 1.825 | 0.034 |
| ENSDARG00000015134 | *camkk1b* | Calcium/calmodulin-dependent protein kinase kinase 1b | -1.277 | 0.023 | -1.594 | 0.034 |
| ENSDARG00000063333 | *nup210* | Nucleoporin 210 | 0.633 | 0.043 | 1.14 | 0.038 |
| ENSDARG00000073699 | *col9a1a* | Collagen, type IX, alpha 1a | 1.014 | 0.029 | 1.483 | 0.038 |
| ENSDARG00000027963 | *camkv* | CaM kinase like vesicle associated | -0.744 | 0.022 | -0.764 | 0.038 |
| ENSDARG00000074443 | *gas7a* | Growth arrest-specific 7a | -0.524 | 0.035 | -0.832 | 0.038 |
| ENSDARG00000053453 | *mpp2a* | MAGUK p55 subfamily member 2a | -0.797 | 0.023 | -0.976 | 0.038 |
| ENSDARG00000078475 | *klhl23* | Kelch like family member 23 | -1.064 | 0.018 | -1.011 | 0.038 |
| ENSDARG00000096319 | *-* | Uncharacterized protein | -0.906 | 0.022 | -0.98 | 0.038 |
| ENSDARG00000056262 | *slc35g2a* | Solute carrier family 35 member G2a | -0.661 | 0.023 | -0.815 | 0.038 |
| ENSDARG00000036005 | *pitpnb* | Phosphatidylinositol transfer protein beta | 1.022 | 0.022 | 1.2 | 0.039 |
| ENSDARG00000058094 | *ciarta* | Circadian associated repressor of transcription a | -0.844 | 0.022 | -0.93 | 0.039 |
| ENSDARG00000101980 | *bmper* | BMP binding endothelial regulator | 1.225 | 0.022 | 1.375 | 0.04 |
| ENSDARG00000007382 | *ubtd1a* | Ubiquitin domain containing 1a | -1.338 | 0.022 | -1.397 | 0.04 |
| ENSDARG00000057241 | *tnfsf10* | Tumor necrosis factor superfamily member 10 | 0.927 | 0.027 | 1.327 | 0.04 |
| ENSDARG0000010958 | *glmnb* | Glomulin, b | -2.583 | 0.018 | -2.243 | 0.041 |
| ENSDARG00000100000 | *-* | Uncharacterized protein | -0.982 | 0.018 | -0.799 | 0.044 |
| ENSDARG00000036493 | *-* | Uncharacterized protein | 0.622 | 0.031 | 0.932 | 0.044 |
| ENSDARG00000086172 | *acvr1c* | Activin A receptor type 1C | -1.282 | 0.02 | -1.204 | 0.044 |
